# High-Resolution Tandem Mass Spectrometry Indicates Rubisco Activase is Associated with PS I-LHC I-LHC II Membranes

**DOI:** 10.1101/2022.02.13.480267

**Authors:** Ravindra S. Kale, Larry Sallans, Laurie K. Frankel, Terry M. Bricker

## Abstract

During proteomic investigations examining the mobile LHC II chlorophyll antenna complex associated with Photosystem I (PS I), we identified, with very high confidence, the association of Rubisco Activase with PS I-LHC I-LHC II membranes. Using very rigorous criteria (p-values ≤ 10^-5^), a total of fifty-three high-quality Rubisco Activase peptides were identified by high resolution tandem mass spectrometry in two biological replicates, each digested with either trypsin or chymotrypsin, independently. Using these criteria, and searching the entire spinach proteome, only proteins previously known to be associated with PS I (the PS I subunits PsaD and PsaL, Lhca1-4, Lhcb1-3), the monomeric Lhcb proteins, Lhcb4-6 (which will be discussed elsewhere), and Rubisco Activase were identified. The presence of Rubisco Activase closely associated with PS I has important implications with respect to the activation of this enzyme and, consequently, the Calvin cycle by PS I, which will be discussed.

## Introduction

Recently, using a styrene-maleic acid copolymer, we have isolated PS I-LHC I-LHC II membranes (Fig. 1) which are highly enriched in the PS I subunits, the LHC I antenna (Lhca1-4) and trimeric LHC II antenna proteins (Lhcb1-3) while being highly depleted of PS II subunits, the cytochrome *b*_*6*_*/f* complex and CF_1_-CF_o_ (1). This preparation appears to represent membrane fragments derived from the chloroplast stroma lamella and/or grana margins (2). The PS I associated with these membranes has been shown to be functionally associated with 1-5 LHC II trimers (1, 3). To further characterize the light-harvesting antennae components present, we have performed a proteomics study of the proteins with apparent molecular masses between ∼20-40 kDa as determined by LiDS-PAGE. The principal results of these studies, with respect to the light-harvesting antenna of PS I, will be presented elsewhere (4).

**Figure 1.**
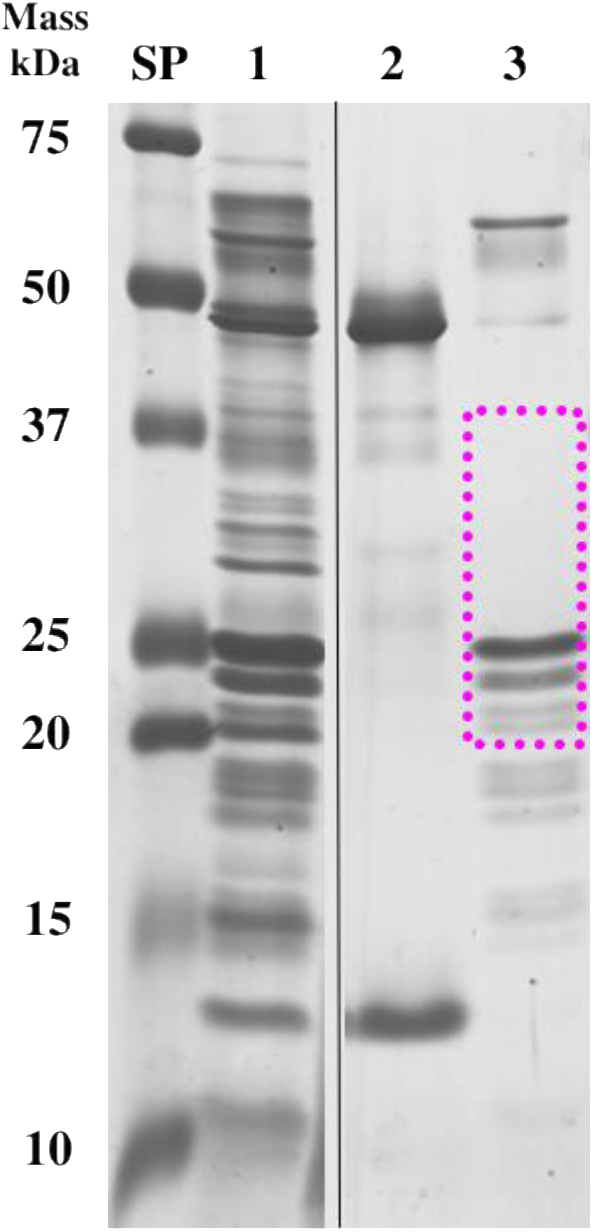
Coomassie-Stained LiDS-PAGE Analysis of Thylakoids, Rubisco and PS I-LHC I-LHC II Membranes. SP, standard proteins, Lane 1, thylakoids, lane 2, sucrose-density purified Rubisco, lane 3, PS I-LHC I-LHC II membranes. The region excised for mass spectrometry analysis (∼20-40 kDa) is outlined by a magenta box. Apparent molecular masses of the standard proteins are shown to the left.

One completely unexpected finding from these experiments was the identification, by high-resolution tandem mass spectrometry, of Rubisco Activase (RA) in association with the PS I-LHC I-LHC II membranes. RA modulates the carbon fixation activity of ribulose 1,5-bisphosphate (RuBP) carboxylase/oxygenase (Rubisco) by removing inhibitory sugar phosphates. The activity of RA is regulated by both the ATP/ADP ratio and by light (see (5) for review).

## Materials and Methods

PS I-LHC I-LHC II-enriched membranes were prepared as previously described (1, 6). LiDS-PAGE (Lithium Dodecyl Sulfate polyacrylamide gel electrophoresis) (7) was performed using a non-oxidizing gel system (8). Electrophoresis was stopped when the proteins entered the resolving gel ∼1.0 cm, the gel was stained with Coomassie blue, de-stained, and a large protein band, which in standard-length gels would represent a molecular mass range of about 20-40 kDa (Fig. 1), was excised. Standard “in gel” digestion protocols were used for both trypsin and chymotrypsin proteolysis. These experiments were performed in duplicate.

For mass spectrometry, the proteolytic peptides were isolated from the gel fragments using standard methods and resolved by high performance liquid chromatography on a C:18 reversed-phase column. Chromatography conditions were as described in (9); data were collected during the acetonitrile gradient and during the wash phase (80% acetonitrile) of the C:18 column. Electrospray ionization was used to introduce the resolved proteolytic peptides into a Thermo Scientific Orbitrap Fusion Lumos mass spectrometer. The samples were analyzed in a data-dependent mode with one Orbitrap MS^1^ scan acquired simultaneously with up to ten linear ion trap MS^2^ scans. The ThermoFisher RAW files were analyzed by the MassMatrix Program (10). A FASTA library containing the entire *Spinacia oleracea* L. (var. Monoe-Viroflay) proteome (11) was searched, as was a decoy library containing the reversed amino acid sequences of the proteome. For a positive protein identification, no hits to this decoy library were allowed (decoy hits = 0.00%). Rigorous p-value thresholds were used for preliminary identification of peptides with a *pp-tag* value of ≤ 10^-5^ and either *pp* or *pp2* being ≤ 10^-5^ (10, 12, 13). The allowed mass error of the precursor MS^1^ ion was required to be ≤ 5.0 ppm. The MS^2^ spectra of the peptides exhibiting these *p*-value and mass accuracy thresholds were then examined manually with the quality of the data being confirmed. Only peptides with charge states of ^+^3 or lower were considered. Finally, for a positive protein identification, at least four peptides meeting the criteria noted above were required in each of the trypsin and chymotrypsin replicate experiments. Typically, in proteomics experiments, proteins are identified using much less stringent criteria. It is very common for p-values of 10^-1.3^ to be applied (14) often with the provision that at least two peptides be identified. We feel that, given the large number of peptides examined in modern Orbitrap-class instruments (>100,000 in our experiments), and the instrument’s high sensitivity, that the use of low stringency criteria can be very misleading.

When applying the stringent criteria used in our experiments, the previously characterized subunits of PS I (PsaD, PsaL and Lhca1-4) (15) and the trimeric LHC II proteins Lhcb1-3 (1-3) were identified. The proteins Lhcb4-6, which had not been previously demonstrated to be associated with PS I were also detected and will be discussed elsewhere (4). The only other protein meeting the stringent identification criteria described above was RA, which had not been previously identified as being associated with PS I.

## Results and Discussion

Fig. 1 illustrates the overall protein composition of PSI-LHC I-LHC II membranes in comparison to thylakoid membranes and sucrose gradient-purified Rubisco. As described earlier (1, 16), the PS I-LHC I-LHC II membranes are highly enriched in the PS I-LHC I supercomplex subunits (PsaA and PsaB, Lhca1-4 and Lhcb1-3, and numerous other lower molecular mass subunits of the supercomplex). Purified Rubisco contains two subunits, the LSU at about 52 kDa and the SSU at about 12 kDa (neither of these were detected in our proteomic experiments). The thylakoid membranes contain these protein components as well as proteins associated with Photosystem II, the cytochrome *b*_*6*_*/f* complex, CF_1_-CF_o_, and numerous other identified and unidentified components. For the mass spectrometry experiments, the region of the polyacrylamide gel represented in Figure 1, lane 3 (PS I-LHC I-LHC II membranes) which encompasses the ∼20 - 40 kDa region (outlined in magenta) was excised, the proteins contained in this region were digested with either trypsin or chymotrypsin, the peptides were then extracted and subjected to high-resolution tandem mass spectrometry.

Figure 2 illustrates the quality of the mass spectra used in this study. The tandem mass spectrometry data collected for the M^+3^ tryptic peptide of RA:^104^K-^123^K are illustrated. The observed mass accuracy for the parent ion was 0.6 ppm. The *pp, pp*_*2*_ and *pp*_*tag*_ values for this peptide were 10^-5.5^, 10^-5.1^ and 10^-5.0^ (10), respectively, and consequently, these were among the lowest quality peptides used in this study. Even this relatively lower quality peptide, however, exhibited nearly complete y- and b-ion series. This allowed near unequivocal identification of the peptide and this result demonstrates that the use of *p* values ≤ 10^-5^ provided extremely high-quality peptide identifications. In total (considering both replicates), 41 tryptic peptides and 12 chymotryptic RA peptides fulfilled these criteria.

**Figure 2.**
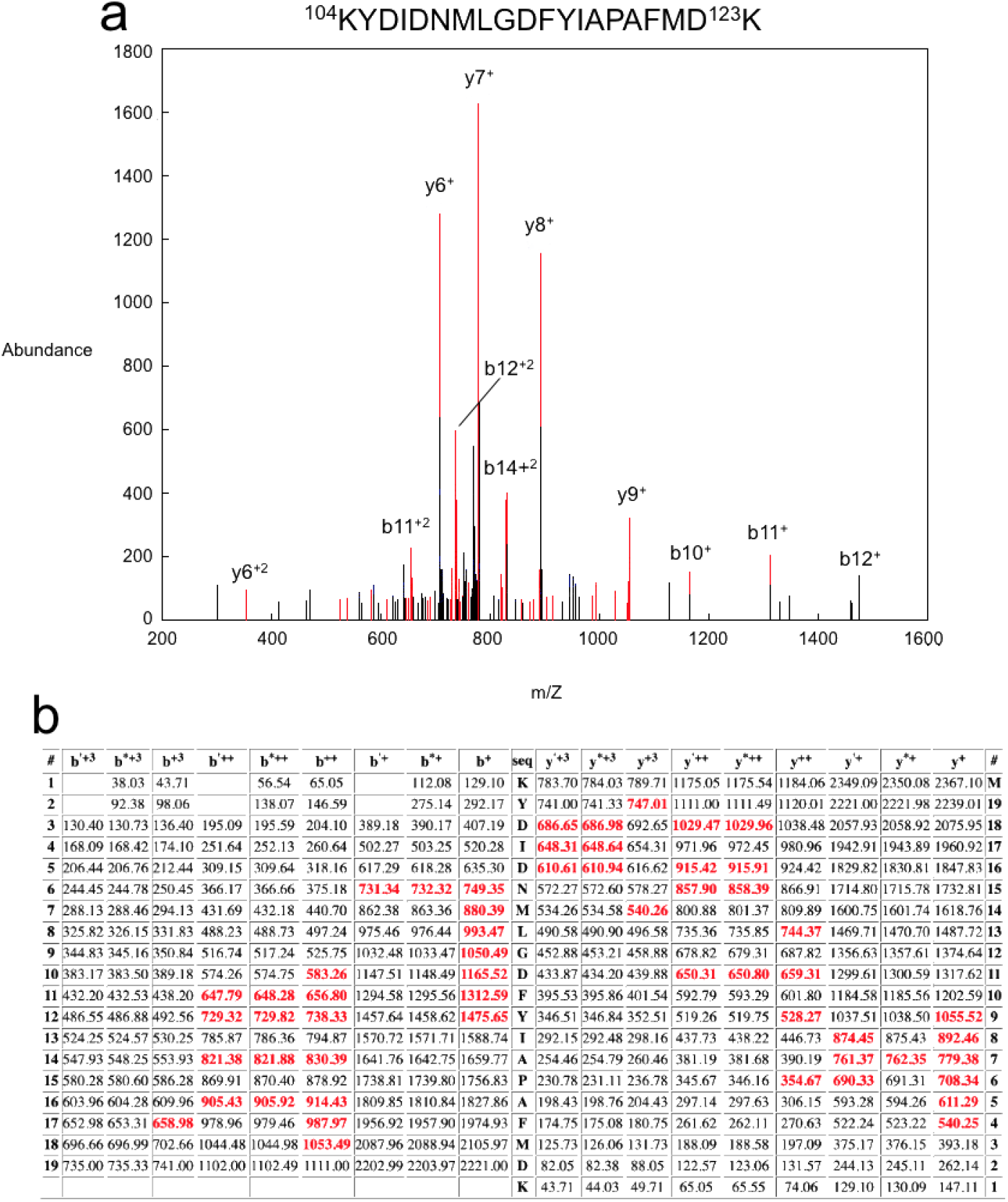
Quality of the Tandem Mass Spectrometry. Shown is the mass spectrometry result for the tryptic peptide of RA:^104^K-^123^K. a, The CID spectrum of the M^+3^ peptide RA:^104^K-^123^K. Identified CID fragment ions are shown as red lines, unidentified ions are shown as black lines. A subset of the identified ions is labeled. b, Table of all predicted masses for the y- and b-ions generated from this peptide sequence. Ions identified in the CID spectrum above are shown in red, those not identified are shown in black. The b’^++^, b’^+^ y’^++^ and y’^+^ ions are generated by the neutral loss of water while the b*^++^, b*^+^ y*^++^ and y*^+^ ions are generated from the loss of ammonia. The *p*-values for this peptide were *pp* = 10^-5.5^, *pp*_*2*_ = 10^-5.1^ and *pp*_*tag*_ = 10^-5.0^ (10). The observed mass accuracy for the parent ion (MS^1^) was -0.6 ppm (predicted mass of peptide = 2367.1036 Da, observed mass of peptide = 2367.1021 Da).

Figure 3 illustrates the mass spectrometry coverage of RA. The results using both trypsin and chymotrypsin are shown. Using trypsin alone, 61% coverage was observed, while using chymotrypsin alone, 43% sequence coverage was observed. In combination, these two proteases achieved an overall sequence coverage of 75%. This is similar to the sequence coverage obtained for the known PS I-LHC I supercomplex subunits (80 ± 11%).

**Figure 3.**
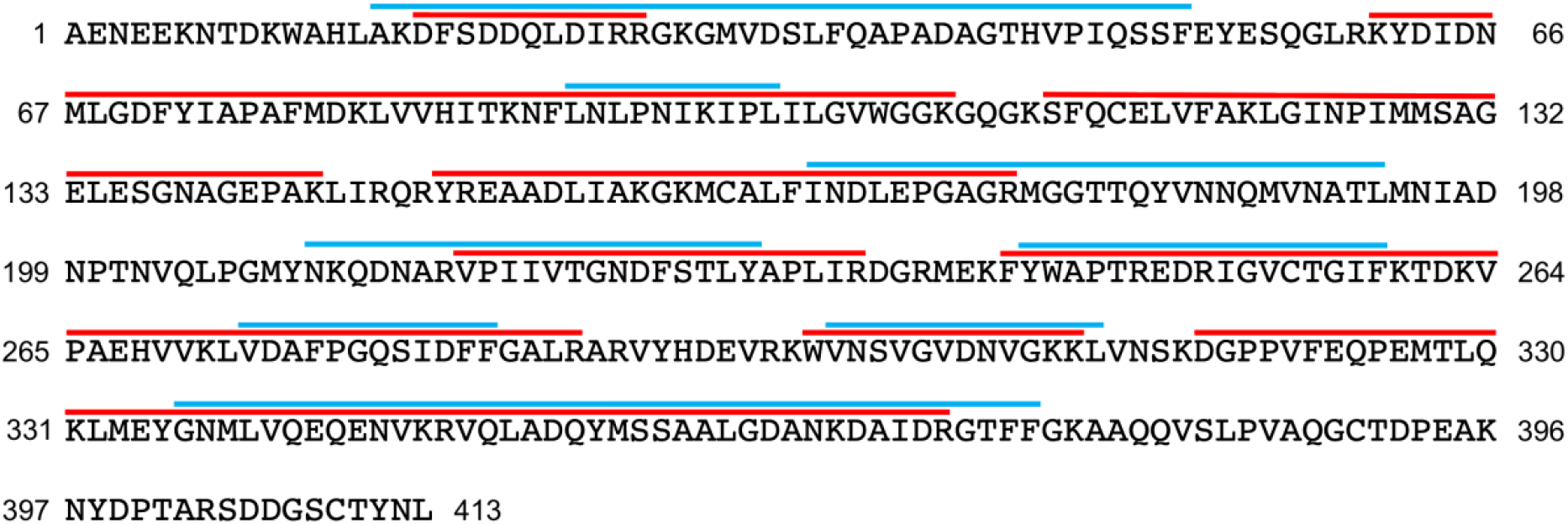
Mass Spectrometry Sequence Coverage of RA. The sequences covered by tryptic peptides are shown by red lines, the sequences covered by chymotryptic peptides are shown by cyan lines. Individual lines may represent overlapping peptides identified by each enzyme. Overall sequence coverage was 75%

Semi-quantitative mass spectrometry, when used for the comparison of soluble and membrane proteins, is extremely difficult and should be viewed with considerable skepticism. Nevertheless, to a very first approximation, spectral counting (17) can provide an estimate of the relative abundance of the components analyzed in these experiments. This is shown in Table 1. The average total spectral counts (trypsin + chymotrypsin spectra) for the six core PS-LHC I subunits (PsaD, PsaL and Lhca1-4) is 24.0 ± 1.2. These subunits are known to be present at a stoichiometry of 1 copy per PS I-LHC I supercomplex (15). These six subunits exhibit a very similar number of spectral counts (range = 19.1 ± 1.0 to 28 ± 5.9). The number of spectral counts for RA is 26 ± 3.8. These results would seem to indicate ∼1 copy of RA per PS I-LHC I supercomplex. However, two factors complicate this interpretation. First, RA is a soluble protein and the six supercomplex subunits are intrinsic membrane proteins. The proteolytic fragments of the intrinsic proteins will tend to be more hydrophobic and will elute from the C:18 chromatography column with less efficiency. They will also tend to ionize with lower efficiency. Both factors will lead to lower spectral count estimates for these components. Secondly, RA is approximately twice the molecular mass of most of the PS I-LHC I subunits (Table 1). Larger proteins, on average, yield a larger number of spectral counts than smaller proteins. Consequently, we estimate that 0.5-1.0 copy of RA may be present per PS I-LHC I supercomplex in these membranes.

**Table I.** Spectral Counts of Identified PS I-LHC I Supercomplex Subunits and RA

|  | <sup>1</sup>Trypsin | <sup>1</sup>Chymotrypsin | Total | Apparent<br><sup>2</sup>Mass (kDa) |
| --- | --- | --- | --- | --- |
| <sup>3</sup>Known PS I-LHC I Subunits: |  |  |  |  |
| PsaD | <sup>4</sup>15.5 ± 0.7 | 8.0 ± 1.4 | 23.5 ± 0.7 | ~20 |
| PsaL | 9.5 ± 0.7 | 9.5 ± 0.7 | 19.0 ± 0.1 | ~15 |
| Lhca1 | 7.5 ± 0.7 | 19.0 ± 1.4 | 26.5 ± 1.5 | ~21-23 |
| Lhca2 | 4.5 ± 0.7 | 16.5 ± 0.7 | 21.0 ± 1.0 | ~21-23 |
| Lhca3 | 7.0 ± 1.4 | 21.0 ± 5.7 | 28.0 ± 0.7 | ~21-23 |
| Lhca4 | 7.0 ± 1.4 | 19.5 ± 2.1 | 26.5 ± 2.1 | ~21-23 |
| RA | 20.5 ± 3.5 | 6.0 ± 1.4 | 26.0 ± 3.8 | ~41-44 |

<sup>1</sup> Average of two biological replicates; all peptides included in the spectral count calculations exhibit pp or pp<sub>2</sub> ≤ 10<sup>-5</sup> and pp<sub>tag</sub> ≤ 10<sup>-5</sup>
<sup>2</sup> From SDS-PAGE
<sup>3</sup> These subunits are present at a stoichiometry of 1 copy per PS I-LHC I supercomplex (19)
<sup>4</sup> Average of two biological replicates; mean ± 1.0 S.D.

As noted in the introduction, light and a high ATP/ADP ratio is required to activate RA. With respect to light-regulation, it was determined that specifically PS I electron transfer was required for the enzyme’s activation (18). Two RA isoforms are present as a result of *trans*-splicing (19). The large isoform of RA contains a C-terminal extension bearing two cystinyl residues (^390^C and ^409^C, spinach numbering, see Fig. 3) which form a disulfide bridge. Reduction of this disulfide by thioredoxin f_red_ leads to this activation of the enzyme (20, 21). Thioredoxin f_ox_ is reduced by ferredoxin_red_ using ferredoxin-thioredoxin reductase, with ferredoxin_ox_ being directly reduced by PS I (for review, see (22)). The activation of RA and, subsequently, Rubisco are critical regulatory steps in the overall redox control of the Calvin-Benson-Bassham cycle (22).

RA is normally considered a soluble protein. Consequently, its redox activation would require diffusion of thioredoxin f_red_ throughout the stromal compartment of the chloroplast. Our identification of RA in association with PS I-LHC I-LHC II membranes raises the interesting possibility that the enzyme may be in close association with PS I, leading to a more direct activation. Such hypothesized “regulatory tunneling” could be analogous to the “substrate tunneling” known to occur in many membrane protein and soluble protein complexes (23). It should be noted that such redox activation would also require that ferredoxin-thioredoxin reductase and thioredoxin also be associated with these membranes. However, their low apparent molecular masses (∼12-13 kDa for both) may have precluded their detection in our experiments. Recently, a spinach supercomplex containing CF_1_-CF_o_ and PS I has been described (24). Interestingly, upon glutaraldehyde crosslinking of the supercomplex, crosslinked products were observed which contained RA, a finding which is completely consistent with our results. The crosslinking partner(s), however, were not identified. In such a supercomplex, both the ATP requirement and the PS I electron transport requirement for RA activation could, hypothetically, be fulfilled.

## Conclusions

High resolution tandem mass spectrometry indicates with high confidence that RA appears to be associated with PS I-LHC I-LHC II membranes. A large number of high quality tryptic and chymotryptic peptides (*p* values ≤ 10^-5^) were identified which yielded an overall sequence coverage of 75%. Comparison of the spectral counts obtained for RA and known subunits of the PS I-LHC I supercomplex indicates that RA is present at a stoichiometry of ∼0.5-1 copy per supercomplex. The association of RA with the PS I-LHC I supercomplex could hypothetically facilitate redox activation of the enzyme.

## Limitations

It must be understood that a direct association of RA with PS I, while implied in this study, is by no means unequivocally shown. More direct evidence, such as protein crosslinking of RA to PS I subunits or single particle TEM and/or Cryo-EM studies resolving RA in intimate association with PS I would be most welcomed. Additionally, the stoichiometry of RA, with respect to the known PS I subunits, appears to be ∼0.5-1:1; however this value is suspect given the difficulties in comparing spectral counts between a soluble protein, RA, and hydrophobic membrane protein components (PsaD, PsaL and Lhca1-4) of the photosystem.

## Acknowledgements

This work was solely supported by the United States Department of Energy, Office of Basic Energy Sciences grant DE-FG02-09ER20310 to TMB and LKF.

