## Supplemental Figure 1 for "High-Resolution Tandem Mass Spectrometry Indicates Rubisco Activase is Associated with PS I-LHC I-LHC II Membranes"

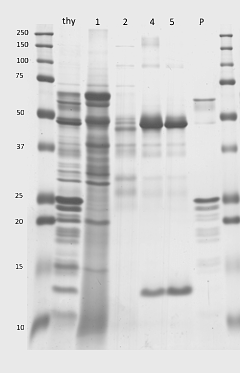

Figure S1. Original Coomassie-Stained LiDS-PAGE Gel used for Figure 1. The left and right most lanes are molecular mass markers which are labeled to the left (apparent mass kDa). Thy, thylakoids, lanes 1-5, sucrose density gradient fractions, P, pellet which constitutes the PS I-LHC I- LHC II membranes (1). Illustrated in Fig. 1 are the molecular mass markers, Thy, lane 5, and P.
